# 3D Cell Nuclear Morphology: Microscopy Imaging Dataset and Voxel-Based Morphometry Classification Results

**DOI:** 10.1101/208207

**Authors:** Alexandr A. Kalinin, Ari Allyn-Feuer, Alex Ade, Gordon-Victor Fon, Walter Meixner, David Dilworth, Jeffrey R. de Wet, Gerald A. Higgins, Gen Zheng, Amy Creekmore, John W. Wiley, James E. Verdone, Robert W. Veltri, Kenneth J. Pienta, Donald S. Coffey, Brian D. Athey, Ivo D. Dinov

## Abstract

Cell deformation is regulated by complex underlying biological mechanisms associated with spatial and temporal morphological changes in the nucleus that are related to cell differentiation, development, proliferation, and disease. Thus, quantitative analysis of changes in size and shape of nuclear structures in 3D microscopic images is important not only for investigating nuclear organization, but also for detecting and treating pathological conditions such as cancer. While many efforts have been made to develop cell and nuclear shape characteristics in 2D or pseudo-3D, several studies have suggested that 3D morphometric measures provide better results for nuclear shape description and discrimination. A few methods have been proposed to classify cell and nuclear morphological phenotypes in 3D, however, there is a lack of publicly available 3D data for the evaluation and comparison of such algorithms. This limitation becomes of great importance when the ability to evaluate different approaches on benchmark data is needed for better dissemination of the current state of the art methods for bioimage analysis. To address this problem, we present a dataset containing two different cell collections, including original 3D microscopic images of cell nuclei and nucleoli. In addition, we perform a baseline evaluation of a number of popular classification algorithms using 2D and 3D voxel-based morphometric measures. To account for batch effects, while enabling calculations of AUROC and AUPR performance metrics, we propose a specific cross-validation scheme that we compare with commonly used k-fold cross-validation. Original and derived imaging data are made publicly available on the project web-page: http://www.socr.umich.edu/projects/3d-cell-morphometry/data.html.

## 1. Introduction

Morphology of a cell nucleus and its compartments is regulated by complex biological mechanisms related to cell differentiation, development, proliferation, and disease [14, 34, 37]. Changes in nuclear morphology are associated with reorganization of chromatin architecture and related to altered functional properties such as gene regulation and expression. Conversely, studies in mechanobiology show that external geometric constraints and mechanical forces that deform the cell nucleus affect chromatin dynamics and gene and pathway activation [32]. Thus, nuclear morphological quantification becomes of major relevance as the studies of the reorganization of the chromatin and DNA architecture in the spatial and temporal framework, known as the 4D nucleome, emerge [11, 36]. Cellular structures of interest in the context of the 4D nucleome include not only the nucleus itself, but also the nucleolus and nucleolar-associating domains, chromosome territories, topologically associating domains, lamina-associating domains, and loop domains in transcription factories [10]. Quantitative analyses of nuclear and nucleolar morphological changes also have medical implications, for example, in detection and treatment of pathological conditions such as cancer [22, 34, 37].

While many efforts have been made to develop cell and nuclear morphological characteristics in 2D or pseudo-3D [12, 25], several studies have suggested that 3D measures provide better results for nuclear morphometry description and discrimination [6, 21]. Although a number of signal processing and computer vision algorithms have been proposed to analyze cell and nuclear morphological phenotypes using 3D representations [8], there is a lack of publicly available 3D cell imaging datasets that could serve for the evaluation of various tools and methods. This limitation becomes of great importance in the modern reality of big data microscopy, when the ability to evaluate different approaches on publicly available data is needed for better dissemination of the current state of the art methods for bioimage analysis [4, 20].

In order to enable objective evaluation of the methods for nuclear morphometric analysis, we create a 3D cell nuclear morphology dataset. The dataset includes 3D fluorescence microscopy volumetric images of cell nuclei and nucleoli of two different cell collections: primary human fibroblast cells and human prostate cancer cells (PC3). In turn, each collection contains images of cells in two different phenotypic states that have previously been shown to exhibit quantifiable changes in nuclear morphology. This allows to evaluate methods for morphological quantification on two binary classification problems.

We also provide a baseline evaluation of simple voxel-based morphometric analysis methods. First, we use automatic segmentation methods to extract individual nuclear and nucleolar binary masks in 3D. We then extract common 2D and 3D voxel-based measures of binary mask morphology and combine them into per-nucleus feature vectors. These feature vectors then used to evaluate a number of machine learning algorithms to provide morphology classification performance baselines. To account for batch effects, while enabling calculations of the Area under the Precision-Recall curve (AUPR) and the Area Under the Receiver Operating Characteristic curve (AUROC) performance metrics, we propose a specific cross-validation (CV) scheme.

To promote the reproducibility of results, facilitate open-scientific development, and enable collaborative validation we will make our workflows, together with underlying source code, documentation, and all derived data from this study available online. Original and derived imaging data are made publicly available on the project web-page: http://www.socr.umich.edu/projects/3d-cell-morphometry/data.html. Additionally, extracted morphometric features are made available for interactive exploration and analysis online via our visual analytics platform SOCRAT [16].

## 2. Dataset preparation

### 2.1. Sample preparation

The dataset is composed of two different cell collections. Each collection includes 3D volumetric images of cells in two phenotypic states that have been shown to exhibit different nuclear and/or nucleolar morphology.

The first collection includes images of primary human fibroblast cells (newborn male) that were purchased from ATCC (BJ Fibroblasts CRL-2522 normal). In order to introduce morphology changes, a part of this collection was subjected to a G0/G1 Serum Starvation Protocol [17]. This protocol is used for cell cycle synchronization and has previously been shown to cause morphology changes in human fibroblasts, affecting nuclear size and shape [31]. As a result, the first collection contains 3D volumetric images of cells in the following phenotypic classes: (1) proliferating fibroblasts (PROLIF), and (2) cell cycle synchronized by the serum-starvation protocol (SS). These classes serve as two categories in a binary morphology classification setting.

The second collection contains images of human prostate cancer cells (PC3). Through the course of progression to metastasis, malignant cancer cells undergo a series of reversible transitions between intermediate phenotypic states bounded by pure epithelium and pure mesenchyme [34]. These transitions in prostate cancer are associated with quantifiable changes in both nuclear and nucleolar structure [22, 35]. Microscope slides of prostate cancer cell line PC3 were cultured in: (1) epithelial (EPI), and (2) mesenchymal transition (EMT) phenotypic states, as described in [35]. Thus, this setting can also be treated as a binary classification task.

### 2.2. Image acquisition

Cells in both collections are labeled with 3 different fluorophores: DAPI (4’,6-diamidino-2-phenylindole), a common stain for the nuclei, fibrillarin antibody (anti-fibrillarin) and ethidium bromide (EtBr), both used for nucleoli staining. Although anti-fibrillarin is a commonly used nucleolar label, we find it to be too specific, which makes the extraction of a shape mask problematic. It has been shown that EtBr can be used for staining dense chromatin, nucleoli, and ribosomes [3]. We find that it provides better overall representation of nucleolar shape. Anti-fibrillarin is combined with EtBr by co-localization to confirm correct detection of nucleoli locations as described below. 3D imaging used a Zeiss LSM 710 laser scanning confocal microscope with a 63x PLAN/Apochromat 1.4NA DIC objective.

**Figure 1.**
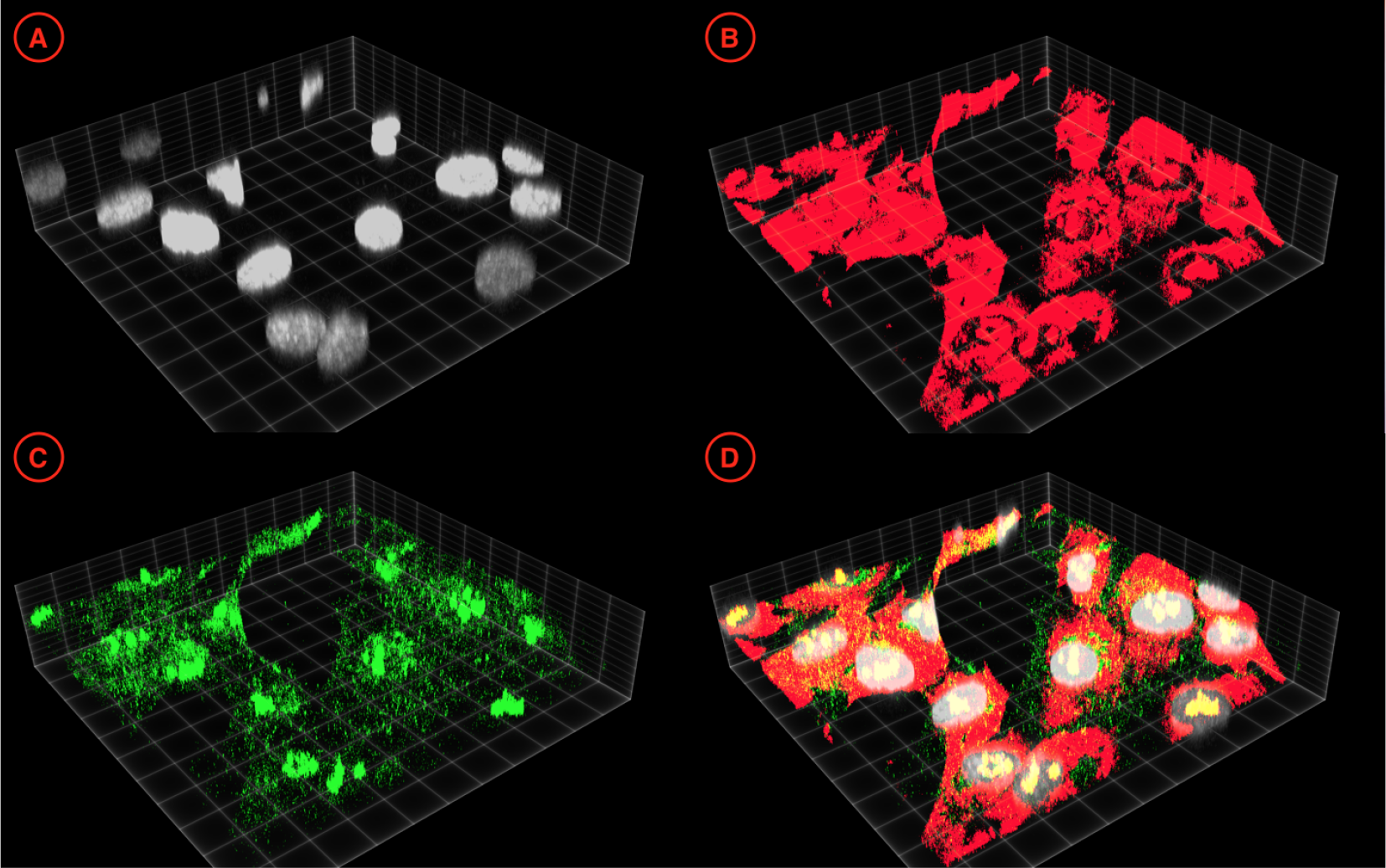
An exemplar 3D visualization of a data sub-volume from the fibroblast cell collection: (A) DAPI channel; (B) EtBr channel;(C) anti-fibrillarin channel; (D) a composite image. Images are thresholded by 25% the for the clarity of visual appearance and visualized using ClearVolume [27].

For multichannel vendor data, the channels are separated and saved as individual volumes labeled as c0, c1, c2, representing the DAPI, anti-fibrillarin, and EtBr channels, respectively, Fig. 1. Each channel-specific volume is then re-sliced into a 1, 024 *×* 1, 024 *× Z* lattice (*Z* = *{30, 50}*), where regional sub-volumes facilitate the alignment with the native tile size of the microscope. All sub-volumes are saved as multi-image 3D TIFF volumes. For every sub-volume, accompanying vendor meta-data are extracted from the original data.

As a result, the fibroblasts collection includes the total of 178 sub-volumes (64 PROLIF and 112 SS), see Table. 1. The PC3 collection includes the total of 101 sub-volumes (50 EPI and 51 EMT), see Table. 2.

**Table 1.** The size of the fibroblast cell collection. Sub-volumes column shows the number of 1024 *×* 1024 *× Z* sub-volumes per channel.

| Class | Sub-volumes | GBs |
| --- | --- | --- |
| PROLIF | 64 | 10.6 |
| SS | 112 | 19.2 |
| TOTAL | 178 | 29.8 |

**Table 2.** The size of the PC3 cell collection. Sub-volumes column shows the number of 1024 *×* 1024 *× Z* sub-volumes per channel.

| Class | Sub-volumes | GBs |
| --- | --- | --- |
| EPI | 50 | 15.7 |
| EMT | 51 | 21.3 |
| TOTAL | 101 | 37.0 |

## 3. Methods

To establish baseline morphometry classification results, we first segment nuclei and nucleoli from the original data sub-volumes. Then, we extract multiple voxel-based morphometric characteristics from 3D binary masks and their 2D projections (2D masks). We use these features to evaluate the performance of a number of widely used classification algorithms. We also assess possible batch effects in data by comparing two different cross-validation techniques.

### 3.1. Nuclear segmentation

Model-based cell segmentation approaches are the most common in bioimage analysis and typically perform well for fluorescence microscopy images of cultured cells [4]. Moreover, they allow to avoid a very labor-intensive process of manual pixel-level expert annotation of large 3D volumetric imaging data. After testing a number of implementations of 3D thresholding-based and watershed-like methods in commonly used bioimage analysis packages, we perform the automatic 3D segmentation of nuclei using Nuclear Segmentation algorithm from the Farsight toolkit [1]. This tool was created specifically to segment DAPI-stained nuclei in 2D or 3D, it does not require a labeled training set, has a convenient command line interface, and demonstrated stable results on these data. The algorithm implements multiple steps which include a graph-cut algorithm to binarize the sub-volumes, a multi-scale Laplacian of Gaussian filter to convert the nuclei to blob masks, fast clustering to delineate the nuclei, and nuclear contour refinement using graph-cuts with alpha-expansions.

After segmentation of the DAPI channel sub-volumes, Fig. 2A, data were converted to 16-bit 3D TIFF files, each segmented nucleus was represented as a binary mask, and given a unique index value. Post-segmentation processing of nuclear masks included 3D hole filling and a filtering step that removed the objects if they span the edge of a tile, are connected to other objects, or their compactness or voxel count values were outside of the empirically estimated interval. This quality control protocol allowed to remove most of the artifacts, as confirmed by visual inspection.

**Figure 2.**
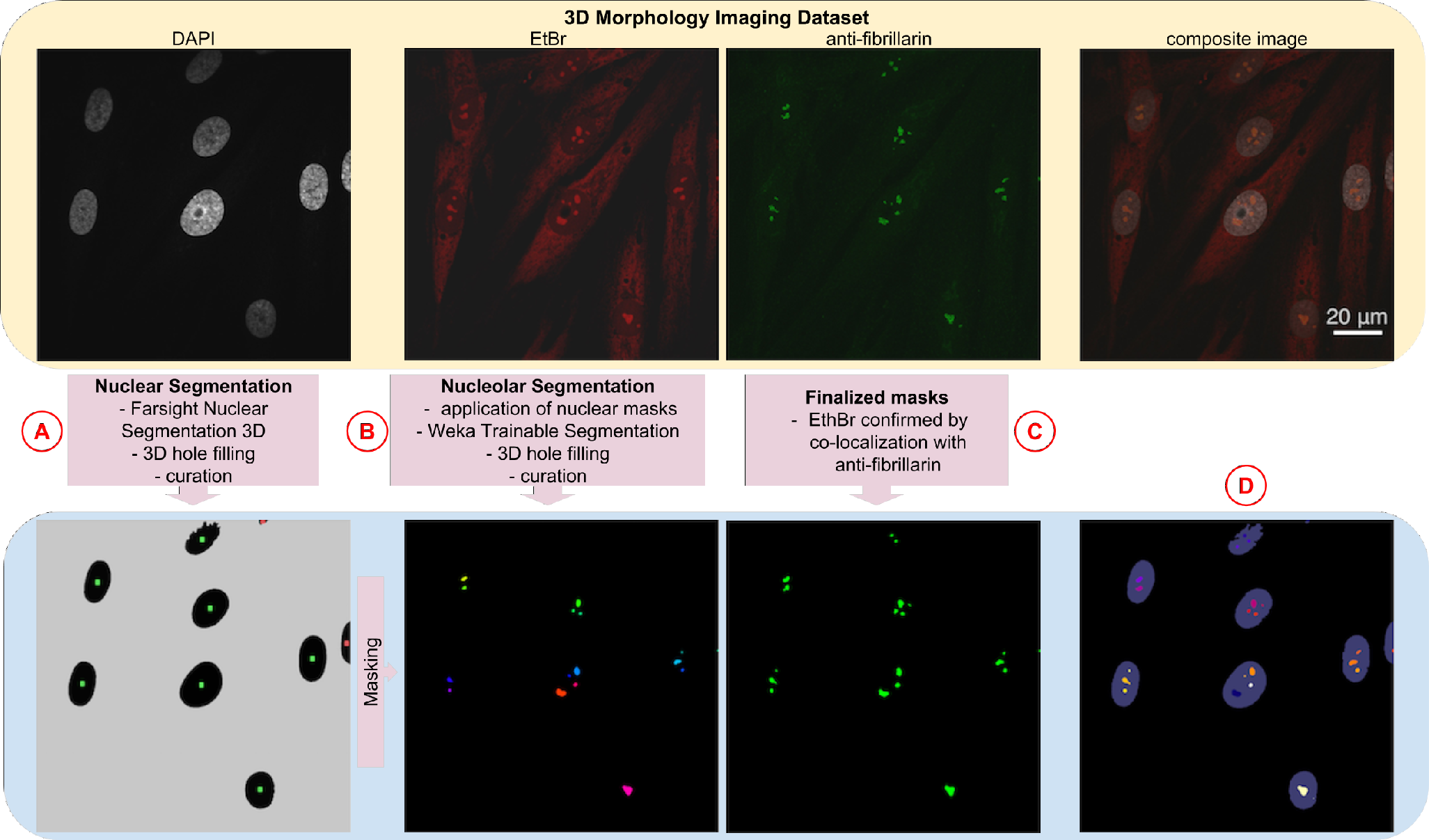
A schematic view of the dataset segmentation protocol and exemplar 2D slices of fibroblast data: (A) steps for the DAPI segmentation process that produces nuclear masks after hole-filling (color-coded by quality control filter); (B) steps for EtBr segmentation that outputs nucleolar masks (colored by connected component labeling); (C) co-localization nucleolar segmented masks with the segmented anti-fibrillarin channel; (D) the composite image of segmented data.

### 3.2. Nucleolar segmentation

Since nucleolar labels are not very specific and produce strong background, see Fig. 1, segmentation of nucleoli using model-based approaches did not demonstrate acceptable results. Therefore, segmentation of objects within the nucleus was performed using the Trainable Weka Segmentation [2], a machine learning tool for microscopy pixel classification bundled with Fiji [29], a commonly used bioimage analysis framework. The Trainable Weka Segmentation plugin is the most popular segmentation tool in ImageJ ecosystem [30], and it is convenient to use for labeling biological structures in 3D images, since it does not require the exact mask contour tracing. Instead, it allows the extraction of a number of features from scarcely labeled pixel groups from both classes, which then are used train a classification algorithm from the WEKA Data Mining software package [9]. Intra-nuclear segmentation was independently performed on EtBr and anti-fibrillarin stained nucleoli. Nuclear masks were used to isolate sub-nuclear segmentations in the EtBr and anti-fibrillarin channels to objects within a nucleus. An individual Random Forest classifier model [18] was created for each channel by using a random selection of 10% of the sub-volumes within that channel for training. Trained models were then applied to all sub-volumes and nucleolar masks were created from the resulting probability maps and labeled as connected components, Fig. 2B. Finally, both EtBr and anti-fibrillarin segmented volumes were used as input to a co-localization algorithm to validate the segmented EtBr-stained nucleoli based on the presence of anti-fibrillarin, Fig. 2C.

The quality control protocol for nucleolar masks was similar to that for the nuclear masks. Since uneven staining can cause occasional segmentation artifacts, filtering step also measured spherical compactness of identified objects [23] and removed the masks if their compactness were outside of the empirically estimated interval.

### 3.3. Voxel-based morphometry

We measure 2D and 3D voxel-based morphometric features of both nuclear and nucleolar binary masks, see Fig. 2D, using image processing library, scikit-image [33].

The 2D feature set includes: area of the object, area of the 2D bounding box, diameter of a circle with the same area as the object, ratio of the object area to the bounding box area, convex hull area, eccentricity, two biggest eigenvalues of the inertia tensor of the region, major and minor axis of an ellipse fitted to the region, the angle between the X-axis and the major axis of the fitted ellipse, perimeter of an object which approximates the contour of the region, the ratio of the region area to the convex hull area.

The set of 3D morphometry features includes: object volume, volume of the 3D bounding box, diameter of a sphere with the same volume as the object, and ratio of the object volume to the bounding box volume.

In oder to aggregate the nucleolar features per nucleus we compute median, minimum, maximum, and standard deviation for each morphometry measure across the nucleoli within one nucleus. Correspondingly, nuclei that do not have any internally positioned nucleoli are excluded from the further analysis. The number of detected nucleoli per nucleus is included as an individual feature. Thus, the total number of features per nucleus is 5 *× N* + 1, where *N* is the number of either 2D or 3D morphometric measures.

We perform exploratory visual analysis of extracted features using SOCRAT [16], a web platform for interactive visual analytics. The goal of visual analytics is to support analytical reasoning and decision making with a combination of highly interactive visualizations and data analysis techniques. SOCRAT implements a visual analytics workflow that encompasses an iterative process, in which data analysts can interactively interrogate extracted morphometric measures in the form of interactive dialogue supported by visualizations and data analysis components. As an example, we include t-Distributed Stochastic Neighbor Embedding (t-SNE) [19] visualizations of both 2D and 3D features generated by SOCRAT [16].

### 3.4. Classification

We compare various supervised classification algorithms from scikit-learn, a popular Python machine learning toolkit [24], including Gaussian Naive Bayes (NB), Linear Discriminant Analysis (LDA), k nearest neighbors classifier (kNN), support vector machines with linear (SVM) and Gaussian kernels (RBF), Random Forest (RF), Extremely Randomized Trees (ET), and Gradient Boosting (GBM). All classifiers use default hyper-parameters. Feature preprocessing includes feature standardization by subtracting the mean and scaling to unit variance of the training set. In this study, we assign the label of the whole image to every single cell extracted from it.

### 3.5. Cross-validation

To evaluate the possible batch effect that could occur during the image acquisition [4], we compare traditional k-fold cross-validation (CV) scheme with the suggested Leave-2-Opposite-Groups-Out (L2OGO) scheme. L2OGO ensures that: (1) all masks derived from one image fall either in the training or testing set, and (2) testing set always contains masks from 2 images of different classes. Unlike Leave-One-Group-Out CV, L2OGO enables per-split evaluation of performance metrics such as the Area under the Precision-Recall curve (AUPR) and the Area Under the Receiver Operating Characteristic curve (AUROC). Since original volumes are of different size and contain different number of nuclei, we joined smaller volumes into bigger groups to reduce class imbalance in testing sets and the variance of the performance metric estimates. Given the class imbalance in L2OGO, we compute AUC, AUPR, and F1 score to compare algorithms [28].

## 4. Results

### 4.1. Fibroblast cells morphometric analysis

After the curation process and the exclusion of nuclei without detected nucleoli, the full collection of segmented fibroblasts consists of total 965 nuclear (498 SS and 470 PROLIF) and 2,181 nucleolar (1,151 SS and 1,030 PROLIF) binary masks. 2D and 3D morphometric measures of nuclear and nucleolar masks are merged into per-nucleus feature vectors as described above. To assess the variability of data, we use t-SNE [19], a dimensionality reduction technique that is well suited for the visualization of high-dimensional datasets on the 2D space. Fig. 3 suggests that the projection of 3D morphometric measures provides better separability of nuclei clusters in the feature space, although still not perfect.

**Figure 3.**
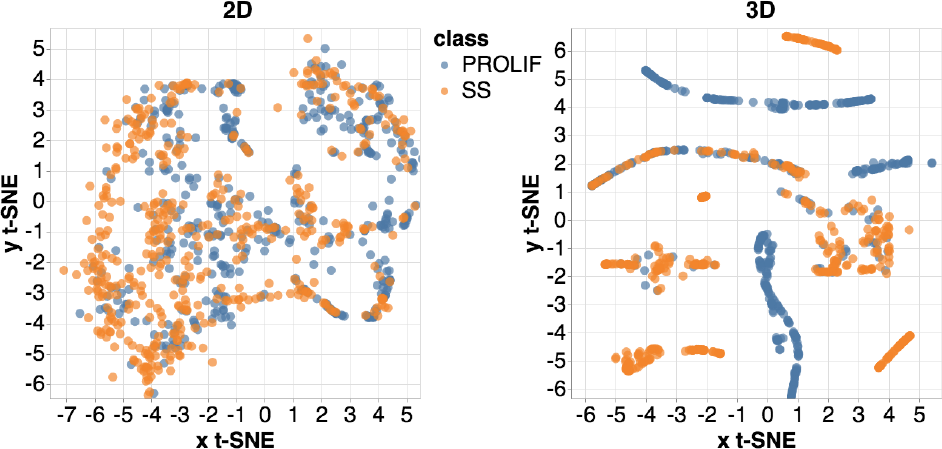
A t-SNE projection of 2D and 3D voxel-based morpho-metric features extracted from binary masks of the fibroblast data. Visualized using SOCRAT [16].

Next, we evaluate the performance of algorithms for Fibroblast morphometric classification on 2 different CV schemes: 20 splits in L2OGO and a 7 times repeated 4-fold CV. Results in Fig. 4 do not show any apparent batch effects in the 2D classification setting. However, 3D performance estimates for all classifiers using L2OGO are more pessimistic compared to 4-fold CV, which indicates the possibility of batch effects and overly optimistic classification results in 4-fold CV. As expected, L2OGO led to a large variance of metrics, especially in the F1 score, which can be explained by classifiers’ sensitivity to different class imbalances in each iteration of this scheme. Within L2OGO, a number of algorithms showed higher performance on 3D morphometry compared to 2D features. The best overall result is achieved by the Gaussian SVM (RBF) classifier in 3D with the median *AUC* = 0.814 *±* 0.245, *AUP R* = 0.724 *±* 0.206, and *F* 1 = 0.709 *±* 0.185).

**Figure 4.**
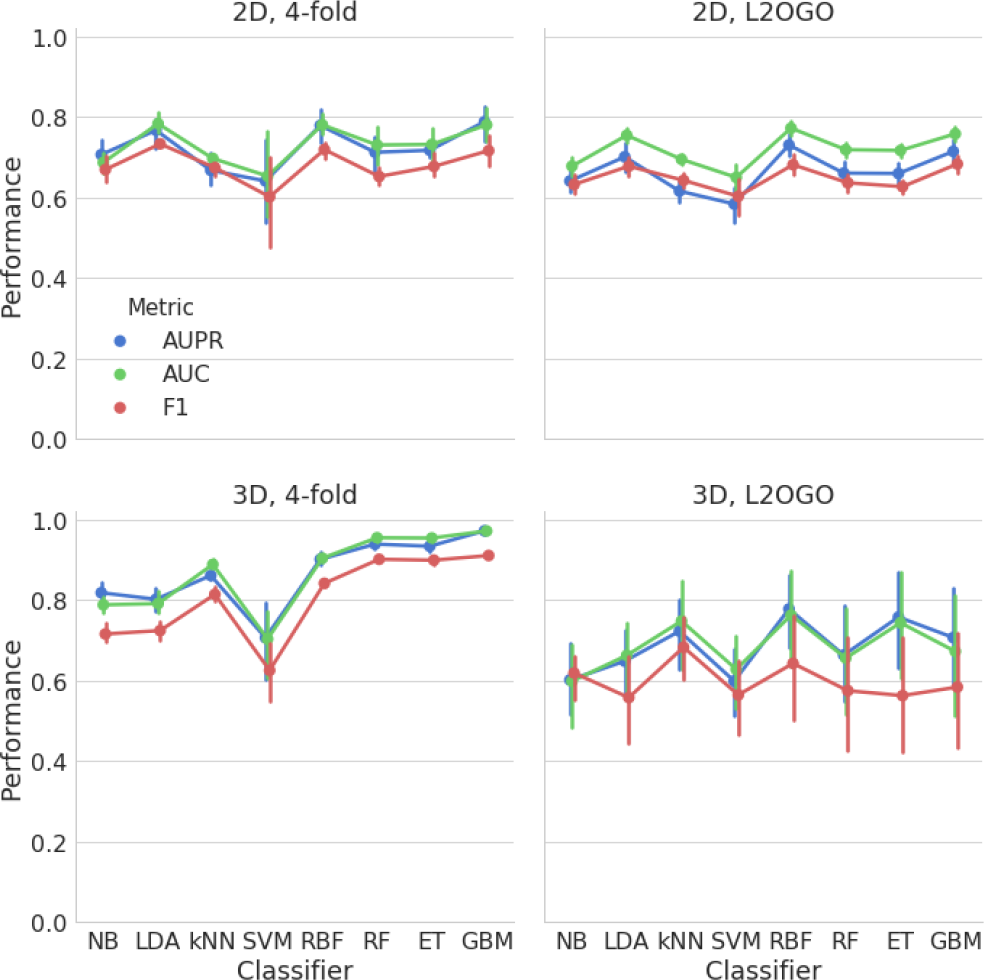
The comparison of cross-validation strategies and commonly used algorithms to evaluate the classification performance and possible batch effects using combined morphometric features of 2D and 3D fibroblast nuclear and nucleolar binary masks.

### 4.2. PC3 cells morphometric analysis

After the exclusion of nuclei without detected nucleoli, the segmented PC3 collection consists of 458 nuclear (310 EPI and 148 EMT) and 1,101 nucleolar (649 EPI and 452 EMT) binary masks. Fig. 5 shows t-SNE projection, demonstrating better cluster separation produced from 3D morphometric measure space, suggesting that 3D feature representations are more informative compared to their 2D counterparts.

**Figure 5.**
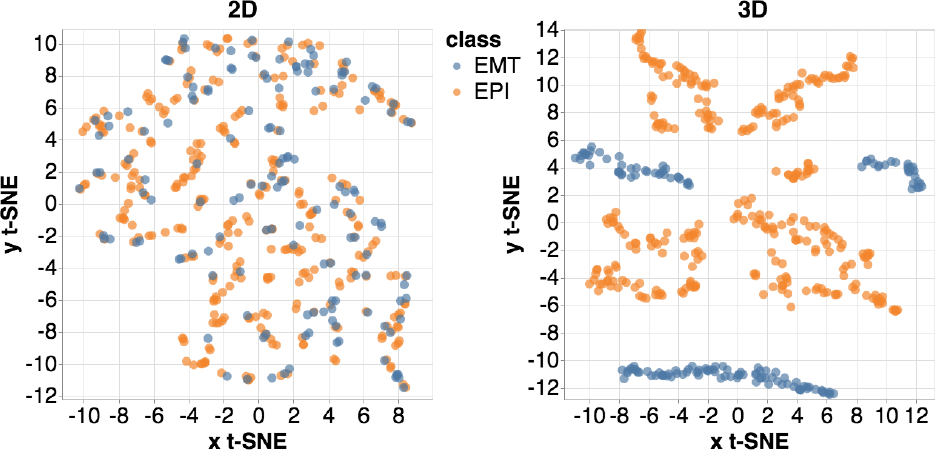
A t-SNE projection of 2D and 3D voxel-based morpho-metric features extracted from binary masks of the PC3 data. Visualized using SOCRAT [16].

After merging smaller EMT groups, L2OGO scheme produced 4 pairs of groups as training and testing sets. Given smaller number of volumes and apparent class imbalance, we compared L2OGO to 4-fold CV repeated 2 times. Similar to the previous experiment, 2D morphometry classification performance was quite similar for both CV schemes, see Fig. 6. However, in 3D, the performance of algorithms degraded as measured by L2OGO CV, such that no methods performed better than in 2D. This can indicate possible batch effects, given the perfect performance estimates for 3 classifiers on 2D features. However, it is hard to judge given the large performance metrics’ variation in 3D. In this case, the best classification by single classifier was the result of applying the Gradient Boosting classifier (GB) with the median *AUC* = 0.774 *±* 0.017, *AUP R* = 0.875 *±* 0.019, *F* 1 = 0.818 *±* 0.018.

Results of classification on both collections suggest that the combination of the voxel-based morphometry and common algorithms with default parameters can provide a good baseline performance. Using 3D masks can improve the performance as it did in Fibroblast classification. However, it suggests that having the three-dimensional information sometimes can lead to more apparent batch effects and, thus, require more complex validation schemes.

**Figure 6.**
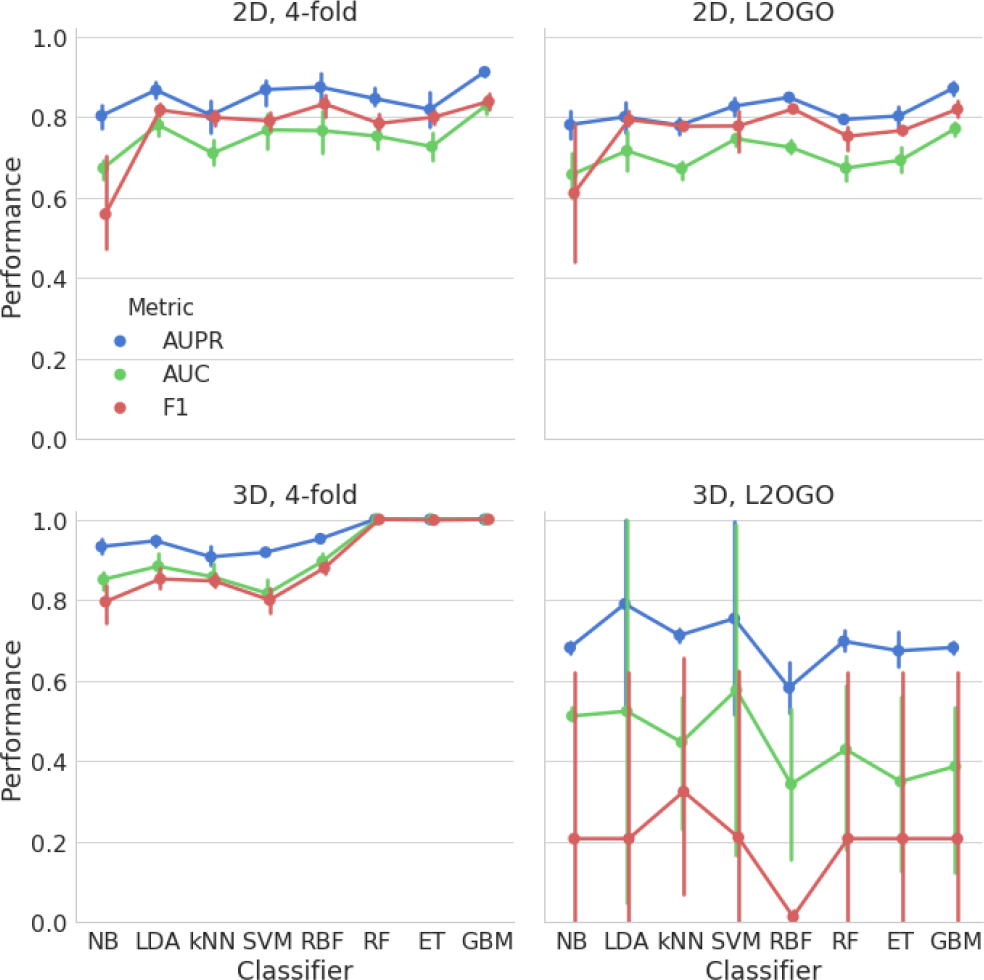
The comparison of cross-validation strategies and commonly used algorithms to evaluate the classification performance and possible batch effects using combined morphometric features of 2D and 3D PC3 nuclear and nucleolar binary masks.

## 5. Discussion

A lack of publicly available 3D cell imaging datasets limits the evaluation of various 3D cell and nuclear morphology analysis solutions. To address this limitation, we present a new dataset that consists of two collections of 3D volumetric microscopic images. Each collection includes images of cells in two phenotypic states and, thus, poses a binary classification problem that can be used for the assessment of cell nuclear and nucleolar morphometry analysis methods. We share these data publicly to promote results reproducibility, facilitate open-scientific development, and enable collaborative validation. To the best of our knowledge, this 3D imaging dataset is one of the largest publicly available datasets of its type.

In order to establish baseline evaluation of simple voxel-based morphometric analysis methods, we provide an example of 3D image processing workflow: from segmentation, to feature extraction, to morphometric analysis. First, we use both model-based and machine learning segmentation methods to extract individual nuclear and nucleolar binary masks in 3D. Then, we extract commonly used 2D and 3D voxel-based measures of binary mask morphology and combine them into per-nucleus feature vectors. Variability of extracted measures between classes is demonstrated via t-SNE projection visualizations. We compare a number of commonly used machine learning classification algorithms on both collections of data using voxel-based morphometric measures. To account for batch effects, while enabling calculations of AUROC and AUPR performance metrics, we also propose a specific cross-validation scheme (L2OGO). Our results indicate potential usefulness of 3D cell imaging data for morphology analysis. However, they also indicate the possibility of stronger batch effects compared to the 2D setting.

As a limitation of this work, the microscope settings did not meet the Nyquist sample rates and may have created distortions in the digitized images [7]. Bigger variability of the performance estimates in 3D using the suggested CV scheme (L2OGO) may be reduced by better class balancing or loss weighting during the each iteration of the cross-validation process. Although produced nuclear and nucleolar binary masks are visually inspected, they are produced by segmentation algorithms rather than hand-labeled by an expert. Thus, these masks should not be considered as a ground truth for segmentation. We provide an example of 3D image processing workflow, which, in general, does not have to always include segmentations step [4]. The size of the produced 3D morphological dataset should be big enough to use segmentation-free deep learning-based morphology analysis approaches [5, 15]. Recent examples in medical image analysis have already demonstrated successful applications of such models in the small data regime [13, 26]. Finally, we assume each cell in the same image to be representative of the same phenotypic label that is provided on the level of the whole image. However, this assumption does not always hold. One 3D volumetric image can contain cells of multiple phenotypes. This can be addressed by using methods for weakly-supervised classification that are robust to label noise.

Imaging protocols, original and segmented data, and the source code are made publicly available on the project web-page: http://www.socr.umich.edu/projects/3d-cell-morphometry/data.html. Additionally, extracted morphometric features are made available for interactive exploration and analysis online via our visual analytics platform SOCRAT [16].

## 6. Conclusion

3D cell microscopy is a powerful technique that enables investigation of biological mechanisms related to morphological changes in cell nucleus through quantitative analysis of changes in its size and shape. The ability to analyze these changes can significantly impact clinical decision-making and fundamental investigation of cell deformation. To our knowledge, we provide the biggest publicly available 3D cell imaging dataset to the date. We describe the data acquisition process and suggest an image processing workflow to establish baseline morphological classification performance. This approach allows an informative evaluation of cell nuclear and nucleolar shapes in the provided imaging data. Public availability of our workflows, source code, documentation, and all derived data from this study facilitates result reproducibility, collaborative method validation, and broad knowledge dissemination in the bioimage analysis community and beyond.

## Acknowledgements

This work was partially supported by the National Institute of Health (grant No. P20 NR015331, U54 EB020406, P50 NS091856, P30 DK089503, P30 AG053760, and T32 GM070449), the National Science Foundation (grant No. 1734853, 1636840, 1416953, 0716055 and 1023115), and the Elsie Andresen Fiske Research Fund.

